# Is reading automatic? Are the ERP correlates of masked priming really lexical?

**DOI:** 10.1101/251884

**Authors:** Dennis Norris, Sachiko Kinoshita, Jane Hall, Richard Henson

**Author notes:** Contact information Dennis Norris, MRC Cognition and Brain Sciences Unit, 15 Chaucer Road, Cambridge, CB2 7EF, United Kingdom.

## Abstract

Humans have an almost unbounded ability to adapt their behaviour to perform different tasks. In the laboratory, this flexibility is sometimes viewed as a nuisance factor that prevents access to the underlying cognitive mechanisms of interest. For example, in order to study “automatic” lexical processing, psycholinguists have used masked priming or evoked potentials to measure “automatic” lexical processing. However, the pattern of masked priming can be radically altered by changing the task. In lexical decision, priming is observed for words but not for nonwords, yet in a same-different matching task, priming is observed for *same* responses but not for *different* responses, regardless of whether the target is a word or a nonword (Norris & Kinoshita, 2008). Here we show that evoked potentials are equally sensitive to the nature of required decision, with the neural activity normally associated with lexical processing being seen for both words and nonwords on *same* trials, and for neither on *different* trials. (150)

---

Human cognition is supremely flexible. Our ability to perform different tasks and make different decisions is almost unbounded. When faced with the same stimulus on different occasions we can choose to process it in many different ways. For example, when presented with a word one may read it aloud, write it down, or count the letters. Each of these actions must necessarily engage at least some different cognitive and neural processes. However, while some processes might change, others might be triggered as a fixed response to the stimulus, regardless of our intentions. We might perhaps have control over how we use the output of those processes, but not over how or whether they operate. Such processes would generally be deemed to be automatic or, in the sense of Fodor (1983), modular. While the distinction between automatic and controlled processing has long been central to research on attention (Atkinson & Shiffrin, 1968; Hasher & Zacks, 1979; Posner & Snyder, 1975), the question of whether a particular mental process is automatic or not is also critical for understanding how to interpret both behavioural and neural measures of cognitive and perceptual processes.

In the study of reading for example, Brown, Gore and Carr (2002) suggested that “Visual word recognition is largely obligatory, in the sense that lexical processing is initiated by the presence of a word in the visual field.” (p. 236). According to that view one might be tempted to assume that the precise task used to investigate word recognition is of little consequence as all tasks will engage the same processes. Their remarks were based on their interpretation of response times in the Stroop task, which is often taken to be a demonstration that word recognition is automatic. However, it has been shown that the Stroop effect relies on the visual conjunction of color and word, and is reduced and may even be eliminated when the two are separated (Risko, Stolz, & Besner, 2005). That is, it is not sufficient for the word to simply be “in the visual field” (Besner, 2001). A related debate concerns whether semantic priming effects are automatic. Several studies have found that priming in the lexical decision task is reduced when the difficulty of the word-nonword discrimination is decreased (Evans, Lambon-Ralph & Woollams, 2012; Shulman & Davidson, 1977; Joordens & Becker, 1997) which implies that priming depends to some extent on the nature of the task (although see Lupker & Pexman, 2010; Milota Widau, McMickell, Juola, & Simpson, 1997). Similarly, Balota and Lorch (1986) found that mediated priming – semantic priming between two otherwise unassociated words which are related indirectly by a mediating word (e.g., lion - (tiger) - stripes) was eliminated when the task was changed from lexical decision to naming. De Wit and Kinoshita (2015) reported that backward-masking the prime to prevent it from reaching conscious awareness eliminated semantic priming when the task was lexical decision, but not when the task was semantic categorization. In sum, in behavioural research “a flexible and adaptive lexical processing system is more consistent with the extant literature” (Yap & Balota, 2015, p.31), and hence considerable effort has been devoted to understanding how cognitive processes are modulated by task demands. There is universal agreement that in order to draw inferences about underlying cognitive processes, we need a good understanding of how experimental tasks are performed. However, it may well be the case that the behavioural consequences of task manipulations reflect changes in the way that the output of fixed and obligatory underlying processes is interrogated: That is, the cognitive processes underlying, say visual word recognition, may be invariant and produce the same range of information, but only the information relevant to performing the specific task is selected. However, common behavioural measures like reaction time (RT) reflect the endpoint in performing a task, and an effect on RT by itself cannot be used to decide at what stage – early or late – task demands modulate cognitive processing. For this reason, researchers have turned to electrophysiological techniques like electroencephalography (EEG), which track brain activity as cognitive processes unfold over time. Perhaps such techniques can more directly reveal the automatic cognitive processes involved in a task such as visual word recognition?

An implicit assumption common to many EEG studies in the field of word recognition is that processing is dominated by a set of core processes. Consequently, the exact choice of task used to study lexical processing is not critically important. Indeed, many EEG studies of visual word recognition have avoided having the participants perform any explicit task at all on the critical trials of interest (e.g. Grainger, Lopez, Eddy, Dufau, & Holcomb, 2011; Holcomb & Grainger 2007, 2009; Massol, Grainger, Dufau & Holcomb, 2010; Massol, Midgley, Holcomb & Grainger, 2011). In those studies, participants were required to detect the occasional occurrence of an animal name while the EEG was measured for non-animal name words. This seems to implicitly assume that the exact choice of reading task is not too important (and also means that there is no behavioural data from the critical trials that can be used to verify that the stimuli are being processed in the same way they are in behavioural experiments designed to examine the same processes).

Other studies have incorporated task manipulations (e.g. Chen, Davis, Pulvermüller & Hauk, 2013; Kiefer & Martens, 2010; Strijkers, Bertrand & Grainger, 2014 ; Vergara-Martínez, Perea, Gómez & Swaab, 2013; Ziegler et al. 1997) and have shown that the nature of the task does modulate the pattern of ERPs. However, a common feature of all of these studies is that the choice of task has only a quantitative influence on either the magnitude or the precise timing of the ERPs. The qualitative pattern of ERPs remains much the same. The general pattern is simply that ERP components associated with lexical processing are attenuated in tasks requiring less linguistic or attentional engagement. These data, generated from a wide range of different laboratory tasks, could therefore be taken to suggest that there is a set of core lexical processes which are invoked in a constant configuration in all tasks, and that tasks differ only in the extent to which they engage those processes.

In contrast to the idea “that lexical processing is initiated by the presence of a word in the visual field” a rather different perspective comes from the idea that perception should be seen as a process of Bayesian inference (e.g. Kersten, Mamassian & Yuille 2004; Knill & Richards, 1996; Norris, 2006; Norris & Kinoshita, 2008). The Bayesian view characterises perception as a process of evaluating competing hypotheses about the state of the world. Different tasks necessarily require the observer to consider different hypotheses, and this should necessarily change behaviour. Perhaps what the data reviewed above are really telling us is that if tasks differ only in the degree of lexical involvement, then this will only be able to effect moderate changes in the size of both behavioural and ERP effects.

According to this view it is also possible that what appear to be lexical effects are actually a consequence of decision processes and not of automatic lexical processing. If this is the case, then perhaps by changing the decision that participants must perform on the stimulus we might be able to effect a much more radical change in the ERPs elicited by words. This would enable us to determine whether ERPs normally elicited by words really are a reflection of automatic lexical processing. Here we investigate this using masked priming.

## The present study

Studies of visual word recognition often use the masked priming procedure introduced by Forster and Davis (1984; see figure 1). As Forster (1998) observed “Masked priming paradigms offer the promise of tapping automatic, strategy-free lexical processing” (p. 203). Masked priming has been used extensively in both the behavioural literature (for a survey, see Kinoshita & Lupker, 2003) and in the EEG literature to study lexical processes in word recognition (for a review of masked priming effects on ERP components, see Grainger & Holcombe, 2009). The most popular view of masked priming is that it provides an “index of lexical access” (Forster, 2004, p. 277). This view is largely based on the observation that, in the lexical decision task, priming is seen for word targets but not for nonword targets. This pattern of priming invites the conclusion that priming is lexical in origin – nonwords do not have representations in the lexicon, so there is nothing to activate, hence there is no priming for nonword targets. This fits well with the view that priming effects are a property of either lexical representations or the structure of the lexicon. If priming operated instead at an orthographic level of processing, one would expect to see equivalent priming for both words and nonwords. The view that nonwords do not show masked priming effects because they do not have a lexical representation is one that is shared by a number of researchers using masked priming as a tool to investigate visual word recognition processes (e.g., Andrews & Hersch, 2010; Davis & Lupker, 2006).

**Figure 1.**
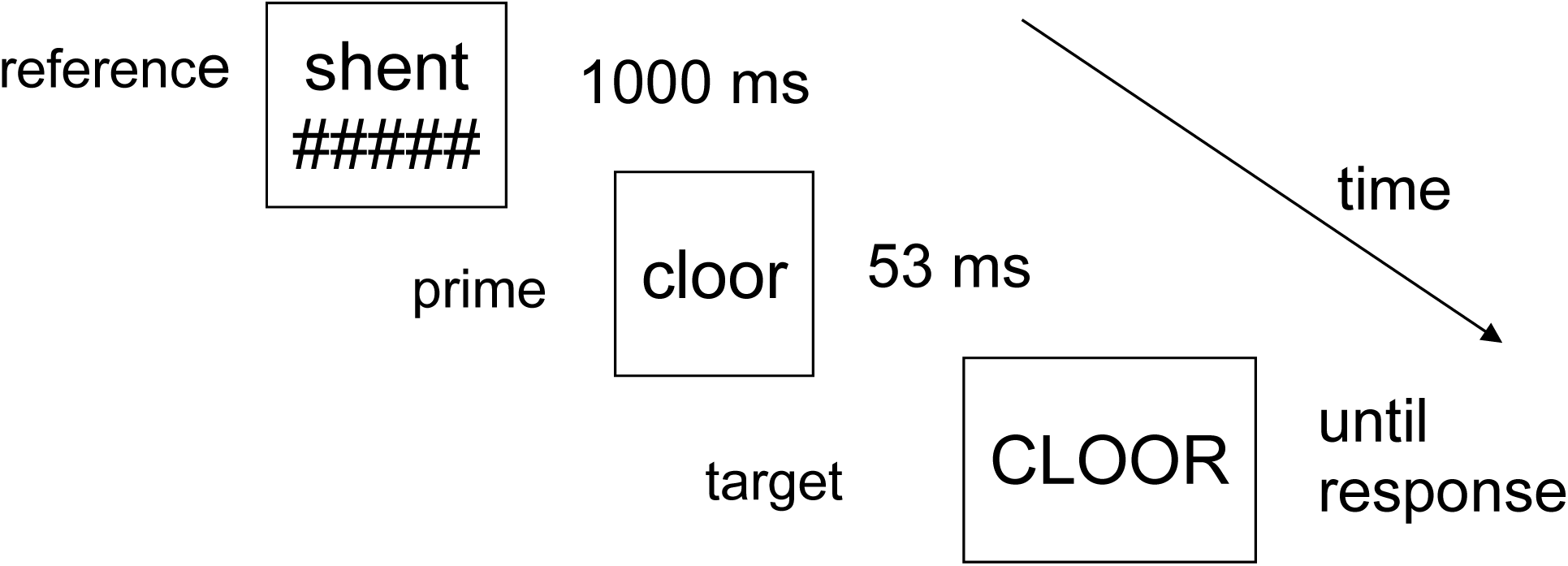
Trial structure. The lexical decision task differs from the same-different task in that the referent is replaced with the word ‘ready’.

More recently, Norris and Kinoshita (2008) have shown that the pattern of masked priming seen in lexical decision can be completely reversed. Under some circumstances it is possible to obtain priming for nonwords while at the same time eliminating priming for words. This pattern of results was predicted from the Bayesian Reader model of word recognition (Norris, 2006, 2009; Norris & Kinoshita, 2012). Norris and Kinoshita used a cross-case, same-different matching task, in which a referent stimulus is presented for 1000ms before the mask and the prime (see Figure 1). The participant’s task is to decide whether the target is the same as the referent or different. When the target was the same as the referent, they found priming for both words and nonwords. When the target was different, there was no priming for either words or nonwords. Furthermore, in the *same* condition, priming effects were equivalent for both words and nonwords. That is, in the same-different task, there is no indication that priming effects are in any way lexically mediated.

In the Bayesian Reader, word recognition is characterised as a process of optimal decision making based on the accumulation of noisy perceptual evidence. It is important to note that here “decision making” does not refer to conscious or strategic decisions, but to the cognitive and perceptual decisions that must underlie all behaviour. One may make a conscious choice to eat an apple rather than a banana, but the process of grasping an apple from a bowl of fruit requires a complex series of perceptual decisions to guide our actions. By decisions we therefore refer to the computations necessary to perform the task, not an optional strategy operating on an invariant, stimulus-driven lexical access process, as assumed by lexical activation models. In the Bayesian Reader, word recognition is modelled as a process of continuous evidence accumulation, and there are no separate stages for “lexical access” and “decision processes” (Norris, 2009). Accordingly, in the Bayesian Reader, the nature of the computation changes as a function of the decision required by the task. In a perceptual identification task, where readers are required to report the word that they have seen, readers must select the word that has the highest posterior probability given the input. In the lexical decision task, the requirement is no longer to identify the best matching word, but to determine whether the input is more likely to be a word than a nonword, and this need not involve extracting sufficient information to be able to uniquely identify the target (for details, see Norris, 2006; 2009; Norris & Kinoshita, 2008; 2012). Norris and Kinoshita suggested that in masked priming, participants treat the mask and prime as a single perceptual object and integrate the evidence^1^ from the prime and target so as to make the perceptual decision required by the task. When the task is changed from lexical decision to same-different, the nature of the decision must necessarily change, and with it the pattern of priming. The full reasoning behind this prediction is given in Norris and Kinoshita (2008), along with a number of simulations. In brief, the basic rationale is that both the lexical decision and the same-different tasks can be characterised as involving a decision as to whether or not the target stimulus is a member of a pre-specified set of stimuli, and that priming only extends to members of that set. In lexical decision this set is the entire lexicon. In the same-different task this set is the single referent stimulus. The model predicts that priming should be obtained for stimuli in that set but not for other stimuli, and this is exactly what was found. In the same-different task there is priming for *same* targets regardless of whether they are words or nonwords, and there is no priming for *different* responses, even when they are words. From the theoretical perspective of the Bayesian Reader, the pattern of data expected in the two tasks is actually the same; there should be priming for items in the set regardless of their lexical status, and no priming for words not in the set. In contrast, from the view that priming is a task-independent function of lexical activation, priming should be seen for words but not nonwords, regardless of task.

An important implication of these results is, therefore, that contrary to the conventional view, priming is not an automatic consequence of the relation between prime and target. The pattern of priming is primarily determined by the nature of the decision/computation required by the task (see Kinoshita & Norris, 2012, for a more general review of task effects in masked priming). If behavioural priming effects are a consequence of the decision required by the task rather than automatic lexical activation, this raises the possibility that the priming effects observed in EEG studies of masked priming might not be specifically lexical either. If this is so, it would mean that priming effects seen in EEG cannot be used to draw conclusions about lexical activation in a task like lexical decision (or even with no task at all), unless we also know how masked priming influences EEG data in the same-different task. If the effects of masked priming on ERPs show the same pattern as the behavioural data, this would imply that these measures cannot be taken as providing evidence about specifically lexical processes either. Here, we will compare masked priming effects on the same word and nonword targets in both the lexical decision task and the same-different task, so as to pit the view that masked priming effects in electrophysiological measures reflect lexical activations against the view that they reflect the evidence contributed by the prime towards the decision required by the task.

Norris and Kinoshita suggested that masked priming effects are entirely a consequence of the decision processes involved in performing the task. If the EEG data, like behavioural data, also reflect the decision processes, we might expect to see ERP priming effects only when we see priming in the behavioural data. For targets requiring a *same* response, even nonwords should produce a pattern like the primed words in lexical decision. For targets requiring a *different* response, words should produce a pattern like nonwords in lexical decision. Alternatively, it is possible that the EEG data – or at least the earliest ERP components that differ between words and nonwords – reflect only the stimulus-driven, task-independent processes. Note that depending on the perspective, the same-different and lexical decision tasks can be seen as producing either a different pattern of results, or exactly the same pattern of results. The behavioural data indicate that lexical decision and same-different differ in terms of whether or not nonwords can be primed, and whether words will always show priming. However, from the theoretical perspective of the Bayesian Reader, the critical predication is really that when the tasks are characterised as making judgements as to whether an item is a member of a task defined set; items in that set can be primed, whereas items not in that set cannot. That is, in this sense, the tasks are the same. The question we are asking therefore is whether pattern of priming observed in the EEG data will be driven by the nature of the stimuli (priming for words not nonwords) or by the decisions required by the task (priming for words or nonwords in the specified set, no priming for either words or nonwords not in the set). Of course, a third possibility is that in the same different task we might see the lexical decision pattern (priming for words but not nonwords), reflecting automatic lexical processes, overlaid by the neural correlates of additional processes involved in performing the same-different task. Note that the same-different task has been used in one pervious ERP study (Muñoz, Perea, García-Orza, & Barber, 2012) however, in that study the stimuli were strings of consonants or symbols so it did not address the question of the automaticity of lexical effects.

Here we investigate these alternative possibilities by directly comparing the lexical decision and same-different tasks while simultaneously recording EEG. The experimental procedure is modelled on that used by Norris and Kinoshita (2008).

## Methods

### Participants

Participants were 32 members of the Cognition and Brain Sciences Unit Volunteer Panel (13 male, mean age 22.7 years). There were 16 participants in each task. All were right-handed native English speakers with normal or corrected to normal vision. Each session lasted for approximately 2 hours, for which participants received an honorarium of £20. The study was approved by Cambridge University Psychological Ethics Committee (reference CPREC 2009.56).

### Procedure

Other than the change in task instructions, the procedure for the lexical-decision and same-different tasks was identical. The trial structure is shown in Figure 1. In the same-different task each trial began with the presentation of a referent stimulus in lowercase letters above a forward mask containing the same number of hash signs as there were letters in the target. The referent and mask were both presented for 1000ms. Both the reference and the forward mask then disappeared; the forward mask was replaced by a prime in lowercase letters presented for 50 ms, which was then replaced by a target presented in uppercase letters. The target remained on the screen until the participant made a response or for 2000ms. In the lexical-decision task the word ‘ready’ appeared in place of the referent stimulus.

Note that, with only a few exceptions (e.g. Carreiras, Vergara, & Perea, 2009; Grainger et al, 2011, exp. 2; Monahan, Fiorentino and Poeppel, 2008, exp. 1), EEG and MEG studies of masked priming have used a different procedure from the behavioural studies and have used a separate backward mask between prime and target. This follows the practice adopted by earlier ERP studies of visual word recognition which directly compared masked vs. unmasked priming. In those studies it was necessary to equate the prime-target SOA while rendering the masked prime unable to be identified. However, the use of a backward mask makes it hard to compare the data with those from behavioural studies that have almost invariably used the Forster and Davis procedure in which the target itself acts as the mask.

In both the lexical decision and the same-different tasks, participants were informed that they would see both real words and nonsense words. In lexical decision, they were told that their task was to decide for each letter string whether it was a word or a nonword, as fast and accurately as possible. Participants were instructed to press the right-hand response button for words and the left-hand response button for nonwords. Participants in the same-different task were told that their task was to decide as fast and accurately as possible whether the second letter string was the same as the first letter string, or different. They were instructed to press the right-hand response button for *same* targets and the left-hand response button for *different* targets. Participants were not informed about the presence of the primes.

Participants were seated in a electromagnetically shielded and sound-proofed room, with low ambient lighting. Stimuli were projected, black-on-white, onto a screen approximately 1.3m in front of the participant, such that they subtended horizontal and vertical visual angles of approximately 1.5° and 0.2° respectively. Stimulus presentation and data collection were performed using E-Prime 2.0 software (Psychology Software Tools, Pittsburgh, PA). Each participant completed 480 test trials that were presented in four blocks, with a self-paced break between the blocks. A different random order of trials was generated for each participant.

### Materials

The critical stimuli used in both tasks were 160 high-frequency words, 160 low-frequency words and 160 nonwords. Half of the items were five letters long, 25% were four letters long, and 25% six letters long. The high-frequency words ranged between 81 and 1,599 occurrences per million (Kucera & Francis, 1967), with a mean of 295. The low-frequency words ranged between 1 and 20 occurrences per million, with a mean of 9. For words, the number of orthographic neighbours (N, as defined by Coltheart, Davelaar, Jonasson & Besner, 1977) ranged between 0 and 18 and the high- and low-frequency words were matched on mean N (4.9 and 4.6, respectively). For the nonwords, N ranged between 0 and 10, with a mean of 2.4. An additional 80 high-frequency words, 80 low-frequency words, and 80 nonwords were selected for use as control primes. Mean frequencies for the high- and low-frequency referent words were 197 per million and 8.7 per million, respectively. The stimuli are listed in the Appendix.

In the same-different task, a further 80 high-frequency words, 80 low-frequency words, and 80 nonwords were used as the referent on trials requiring a *Different* response. Note that the fact that the primes following different reference stimuli were always different form the reference is not a cause for concern. Kinoshita and Norris (2010) have shown that, in contrast to when the prime is visible, masked priming in the same-different task is unaffected by whether the target is predictive of the prime. A similar result is found for lexical decision where Norris and Kinoshita (2008) found that responses were unaffected by whether the unrelated prime had the same lexical status as the target. Both results are as predicted by the Bayesian Reader. Four list versions were constructed; each list contained the same 480 items as targets, and across the four lists, each set appeared in each of the four experimental conditions resulting from a factorial combination of prime type (identity vs. control) and response type (same vs. different). Four participants were assigned to each of the four lists. Participants were given 14 practice items, and there were 2 warm-up items at the start of each test block, which were not included in the analysis.

In the lexical decision task, there were an additional 160 nonwords, half of which had identity primes and half of which had control primes. These extra nonword trials were required in order to equate the number of word and nonword targets. These nonwords were constructed according to the same principles as the critical nonword stimuli. The lexical decision task used the same list counterbalancing procedure as in the same-different task. Four participants were assigned to each of the four lists. Participants were given 14 practice items, and there were 2 warm-up items at the start of each test block.

### EEG recording procedure

The EEG was recorded at the same time as MEG data, from a Neuromag Vectorview system (Elekta, Stockholm, Sweden), but we report only the EEG analyses here because the MEG data show the identical pattern (data available on request). The EEG data were measured using 70 electrodes within an elasticated cap (EasyCap GmbH, Herrsching, Germany), with electrode layout conforming to the extended 10-10% system. A 3D digitizer (Fastrak Polhemus Inc., Colchester, VA, USA) was used to record the locations of the EEG electrodes. The EEG reference electrode was placed on the nose, and the common ground electrode was placed at the left collar bone. Two sets of bipolar electrodes were used to measure vertical and horizontal electro-oculograms (VEOG and HEOG). All EEG electrode impedances were maintained below 10 kΩ.

All further EEG processing was done with SPM5 (www.fil.ion.ucl.ac.uk/spm) and custom scripts written in Matlab (www.mathworks.com). Blink-related artefacts were removed by performing ICA (as implemented in EEGLAB; http://sccn.ucsd.edu/eeglab/), and projecting out those independent components (ICs) whose Pearson correlation with the recorded VEOG exceeded an absolute value of 0.5 and whose Z-score relative to the distribution of Pearson correlations across ICs exceeded 3 (the ICs so identified were then double-checked by manual inspection). Between 1 and 2 such ICs were identified in every run and subject. These corrected data were then downsampled to 100Hz and lowpass filtered to 40Hz. They were epoched from -100ms to +700ms around the onset of each stimulus, allowing for a fixed offset of 34ms (reflecting 2 refreshes of the visual projector at 60Hz), and baseline-corrected from -100ms to 0ms. Bad EEG channels (mean = 2.6, range = 0-5) were detected by a combination of manual inspection and channels in which more than 20% of epochs exceeded an absolute value of 120uV. Epochs in which an incorrect behavioural decision was made, or in which the absolute amplitude of any EEG or HEOG channel exceeded 120uV in a non-bad channel, were discarded (mean of 42.8 and range = 0-102). The epochs were then concatenated across runs, and the EEG data were re-referenced to the average across all non-bad electrodes.

For consistency with other research, all times are measured from presentation of the target. It is important to bear this in mind when comparing masked priming data with that obtained from unmasked stimuli (if, as Norris and Kinoshita claim, the prime and target are integrated and treated as a single perceptual event, then prime onset might be a more appropriate reference point).

### Statistical analysis of EEG data

For statistical analysis, common practice in analysing similar ERP data has been to perform ANOVAs on data averaged across time-windows encompassing the N250 and N400 evoked components (e.g, 150-300ms and 300-550ms respectively). ERPs within that entire range have been attributed to lexical processes (Holcomb & Grainger, 2006). Given that putative lexical priming effects extend throughout this range it is hard to motivate focusing on any particular time windows. Such an approach is particularly unsuited to the study of masked priming. For example, masked priming has been argued to provide a head start in processing (Gomez, Perea & Ratcliff, 2013). In a time-window analysis, the pattern of significance will depend on the exact location of any effect relative to the boundary between time windows. A substantial shift in peak latency within a time-window might not alter the mean amplitude of the signal in that window at all, whereas the same shift in peak latency in the vicinity of a boundary between time windows may produce an interaction with time^2^. Both of these issues are particularly problematic given that the time-windows in which priming effects occur in our same-different task may differ from those in our lexical decision task. Indeed, given that the behavioural priming in the same-different task is so much larger, this seems quite likely to be the case. There are also differences across prior studies in the particular electrodes selected for analysis, and the electrodes showing maximal priming effects in our study may again depend on the task. Given this spatiotemporal uncertainty in how priming effects might manifest themselves in the lexical decision and same-different tasks, we adopted a mass univariate approach in which conditions are contrasted at every single time-point and every single scalp location. The resulting statistics are then adjusted for the multiple comparisons taking into account the spatial and temporal smoothness of the data (see Henson, Mouchlianitis, Matthews, & Kouider, 2008, for a similar approach). This completely avoids the problems associated with making arbitrary decisions as to the time windows or electrodes used in the analysis.

More precisely, we interpolated the EEG data onto a 32 × 32 grid of 3 × 3mm pixels representing the flattened scalp, and tiled these planes for each peristimulus time sample to create a 3-D volume. The time dimension consisted of the 81, 10msec samples in each epoch. The data at each point (voxel) within this volume was subjected to ANOVA with a pooled error term, the nonsphericity of which was estimated by Restricted Maximum Likelihood by pooling over voxels (Friston et al., 2002). The resulting F-statistics for each planned comparison (matching those performed on the behavioural data; see Results) formed a Statistical Parametric Map, which was thresholded at p<.001 uncorrected, and clusters identified in which the number of contiguous voxels was greater than would be expected by chance, as defined by p<.05 family-wise error (FWE) corrected for multiple comparisons using Random Field Theory in SPM12 (Kilner et al., 2005; final estimated smoothness of an equivalent Gaussian FWHM was 24mm × 23mm × 45ms).

## Results

### Behavioural data

The behavioural data for lexical decision and same-different tasks are shown in Tables 1 and 2. The behavioural data replicate the results reported by Norris and Kinoshita (2008).

**Table 1.** Mean Lexical Decision Latencies (RT, in ms) and Percent Error Rates (%E) in Lexical-decision task

|  | Target type |  |  |  |  |  |
| --- | --- | --- | --- | --- | --- | --- |
|  | High-frequency word |  | Low-frequency word |  | Nonword |  |
|  | RT | %E | RT | %E | RT | %E |
| Prime type |  |  |  |  |  |  |
| Identity | 577 | 1.6 | 634 | 5.9 | 711 | 3.8 |
| Control | 615 | 2.4 | 685 | 8.8 | 704 | 3.5 |
| Identity priming effect | 38 | 0.8 | 51 | 2.9 | -7 | -0.3 |

**Table 2.** Mean Decision Latencies (RT, in ms) and Percent Error Rates (%E) in Same-Different matching task.

|  | Target type |  |  |  |  |  |
| --- | --- | --- | --- | --- | --- | --- |
|  | High-frequency |  | Low-frequency |  | Nonword |  |
|  | word |  | word |  |  |  |
| Response type | RT | %E | RT | %E | RT | %E |
| and prime type |  |  |  |  |  |  |
| Same |  |  |  |  |  |  |
| Identity | 473 | 2.3 | 468 | 2.0 | 499 | 3.6 |
| Control | 548 | 7.2 | 548 | 9.1 | 563 | 7.7 |
| Priming effect | 75 | 4.9 | 80 | 7.1 | 64 | 4.1 |
| Different |  |  |  |  |  |  |
| Identity | 569 | 3.9 | 550 | 2.8 | 551 | 4.8 |
| Control | 554 | 4.6 | 563 | 3.9 | 557 | 2.5 |
| Priming effect | -15 | 0.7 | 13 | 1.1 | 6 | -2.3 |

Standard practice in the analysis of masked priming lexical decision experiments is to analyse word and nonword responses separately. In the RT analysis of word responses, there were significant main effects of priming (F(1,15)=95.97, MSE=31551, p < .001) and word frequency (F(1,15)=47.43, MSE=644446, p < .001), and no significant interaction between the two (F(1,15)=2.81, MSE = 754, p=.16). There was no priming for nonwords. In a further analysis combining words (collapsed over frequency) and nonwords, there was a significant interaction between priming and lexical status in RTs (F(1,15)=29.393, MSE=10554, p < .001), and in the errors (F=5.93, MSE=18.46, p < .05).

In contrast, collapsing over frequency in the same-different task, there was no interaction between lexical status and priming (F<1), but a highly significant interaction between priming and response type (same vs. different). This was true of both RTs (F(1,15) = 104.5, MSE = 40952, p < .001) and errors (F(1,15) =23.9, MSE = 275, p<.001). The effect of lexicality was significant in the analysis of RTs (F(1,15)=5.2, MSE = 2354, p < .05) but not errors (F<1), and there was no interaction between lexicality and priming in either RTs (F<1) or errors (F(1,15)=3.5, MSE=46.9, p = .08). The three-way interaction between lexicality priming and same-different was also not significant in RTs (F(1,15)=1.24, MSE=986, p = .28) or errors (F<1). A separate analysis looking only at words revealed no significant effect of word frequency in either RTs (F(1,15) = 1.37, MSE= 1045) or errors (F<1).

In the same-different task, there was priming for *same* items (F(1,15)= 135, p < .001) but not for *different* items (F < 1). Additionally, priming in the same-different task was substantially larger than in lexical decision. This latter result is also predicted by the Bayesian Reader.

### EEG data

#### Grand-mean evoked responses

The general pattern of ERPs can be seen in Figure 2, which shows recordings from electrode P3, where the priming effects were consistently largest in the statistical maps below. Overall, the pattern of ERP data parallels that seen in the behavioural data. Briefly, there are priming effects for words but not nonwords in lexical decision (Figure 2A), particularly high-frequency words (Figure 2B), and for *same* targets (Figure 2C), but not *different* targets (Figure 2D), in the same-different task. Furthermore, the priming effects in the same-different task are much larger and more extensive than in lexical decision.

**Figure 2.**
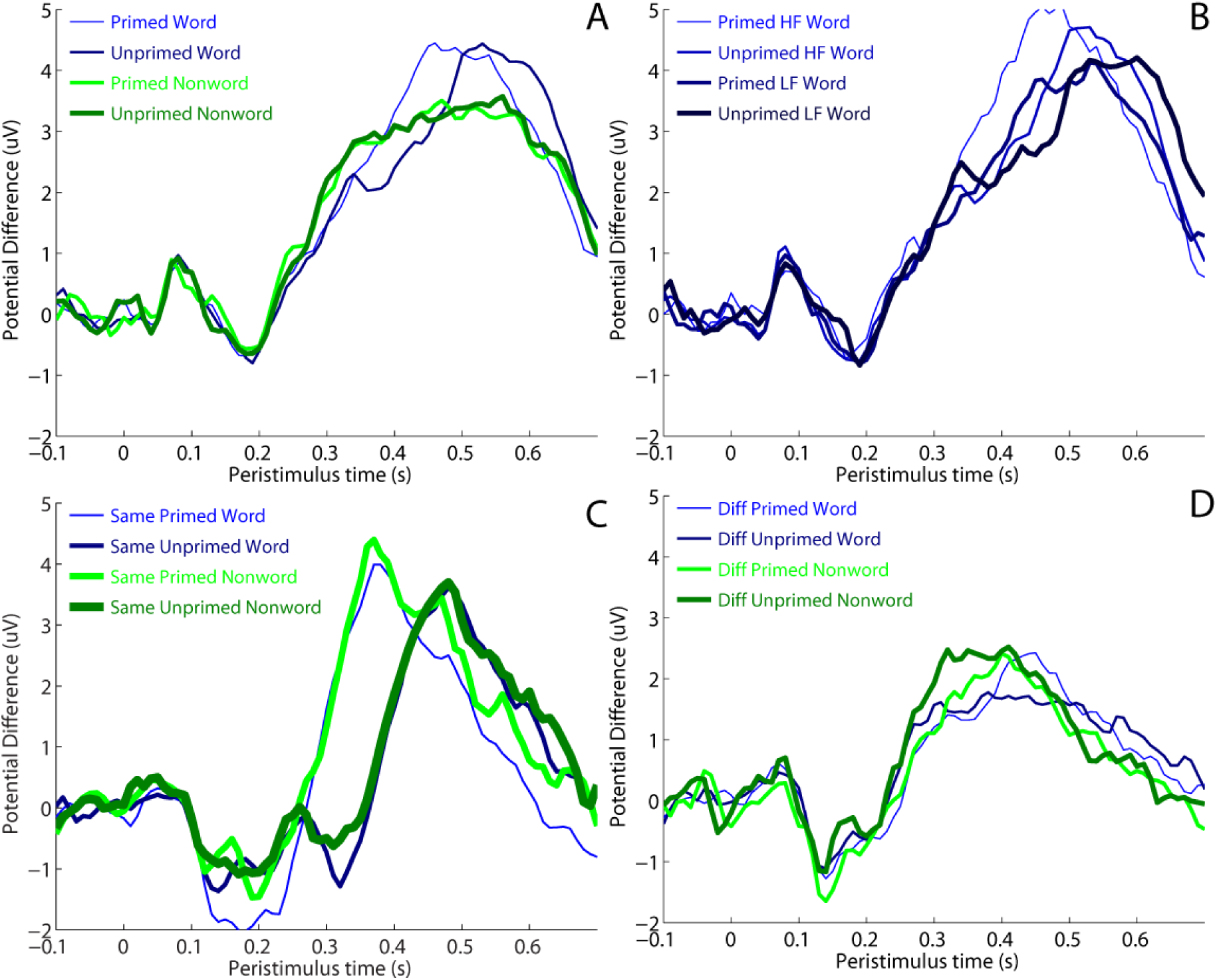
ERPs from Channel P3 (showing maximal priming effects) for (A) primed and unprimed words and nonwords in lexical decision task, (B) primed and unprimed high frequency (HF) and low frequency (LF) words in lexical decision task, (C) primed and unprimed words and nonwords in same trials of same different task, and (D) primed and unprimed words and nonwords in different trials of same different task.

The general pattern of priming in Figure 2 appears to be an acceleration of the entire waveform in both tasks, whereby the ERPs for identity primed trials are shifted earlier in time relative to those for unprimed control trials. This means that there is a crossover between the primed and unprimed ERPs, such that the amplitude of the primed ERP is greater than the unprimed ERP before about 500ms in the lexical decision task, or before 400ms in the same-different task, and then smaller thereafter. With our average reference, this appears as a shift earlier in time of a positive-going component (relative to pre-stimulus baseline) at P3. As a result, the amplitude from 300-500ms is more positive for primed than unprimed words in the lexical decision task, which would correspond to a reduction in a N400 component. There was also evidence of an earlier priming effect on an N200 component over left temporoparietal sites, as detailed below. However, rather than focusing on just these components (whose latency depends on priming), we next tested the significance of condition differences in amplitude at all timepoints and scalp positions.

#### Statistical analysis of priming effects

Figures 3, 4 and 5 show the thresholded space-time SPMs for the three conditions showing priming: words in lexical decision, and both words and nonwords in the *same* condition in same-different task (none of the other priming comparisons, e.g, for nonwords in lexical decision, or for different responses in the same-different task, showed suprathreshold effects).

**Figure 3:**
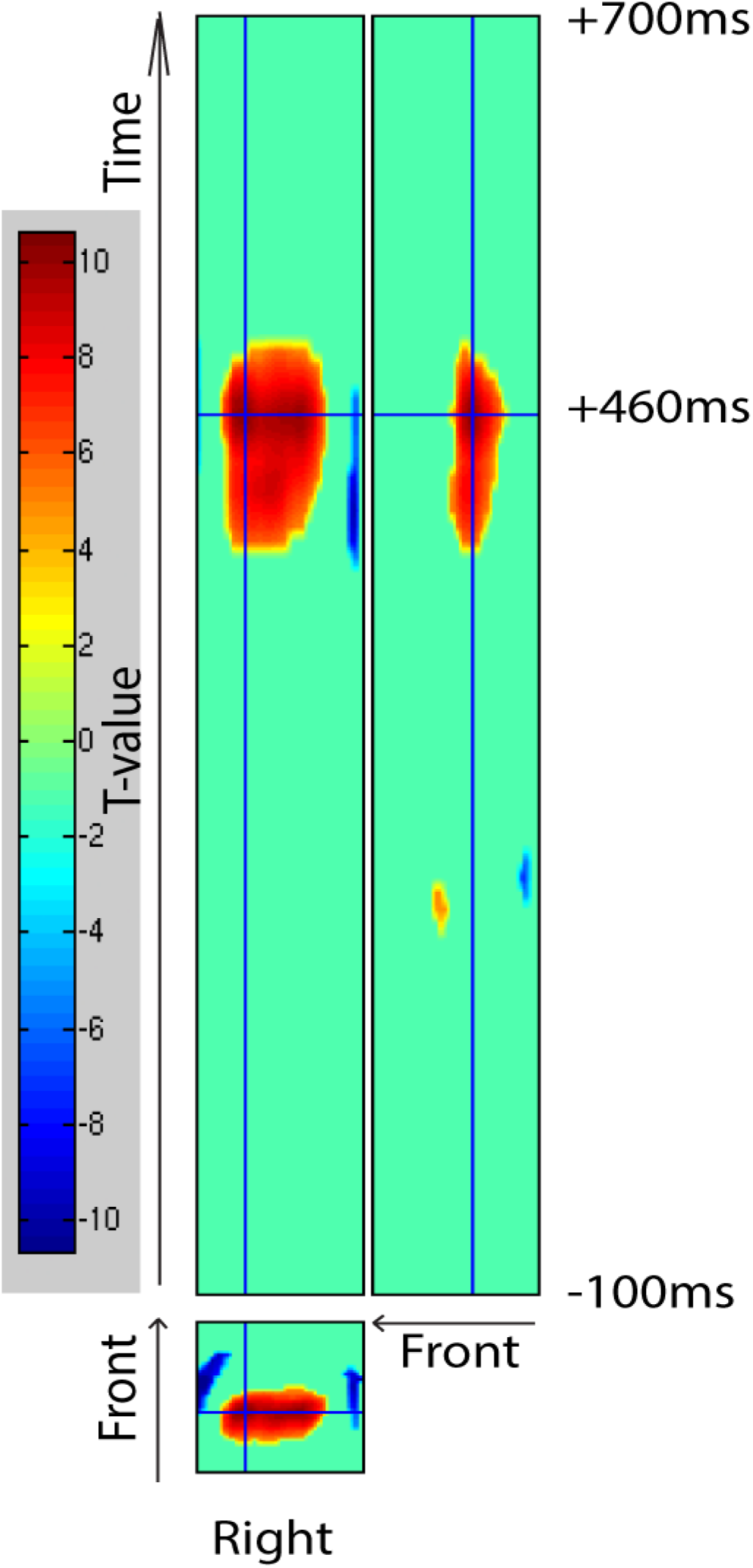
Two sections through the 3D scalp-time Statistical Parametric Map (SPM) for the ERP contrast of primed versus unprimed words in the lexical decision task. The bottom section shows the topography over the scalp at 460ms (relative to an average reference), with the cross-hair centred on the maximum close to electrode P3, while the top section shows a slice through peristimulus time (vertical access) against left-right position at the maximal statistical difference over central sites. The SPM is thresholded such that only clusters of points where the p-value was less than .001 and the probability of obtaining as many such contiguous points by chance was less than .05, corrected for multiple comparisons using random field theory (see Methods). Warm-cold colours indicate positive-negative differences.

**Figure 4:**
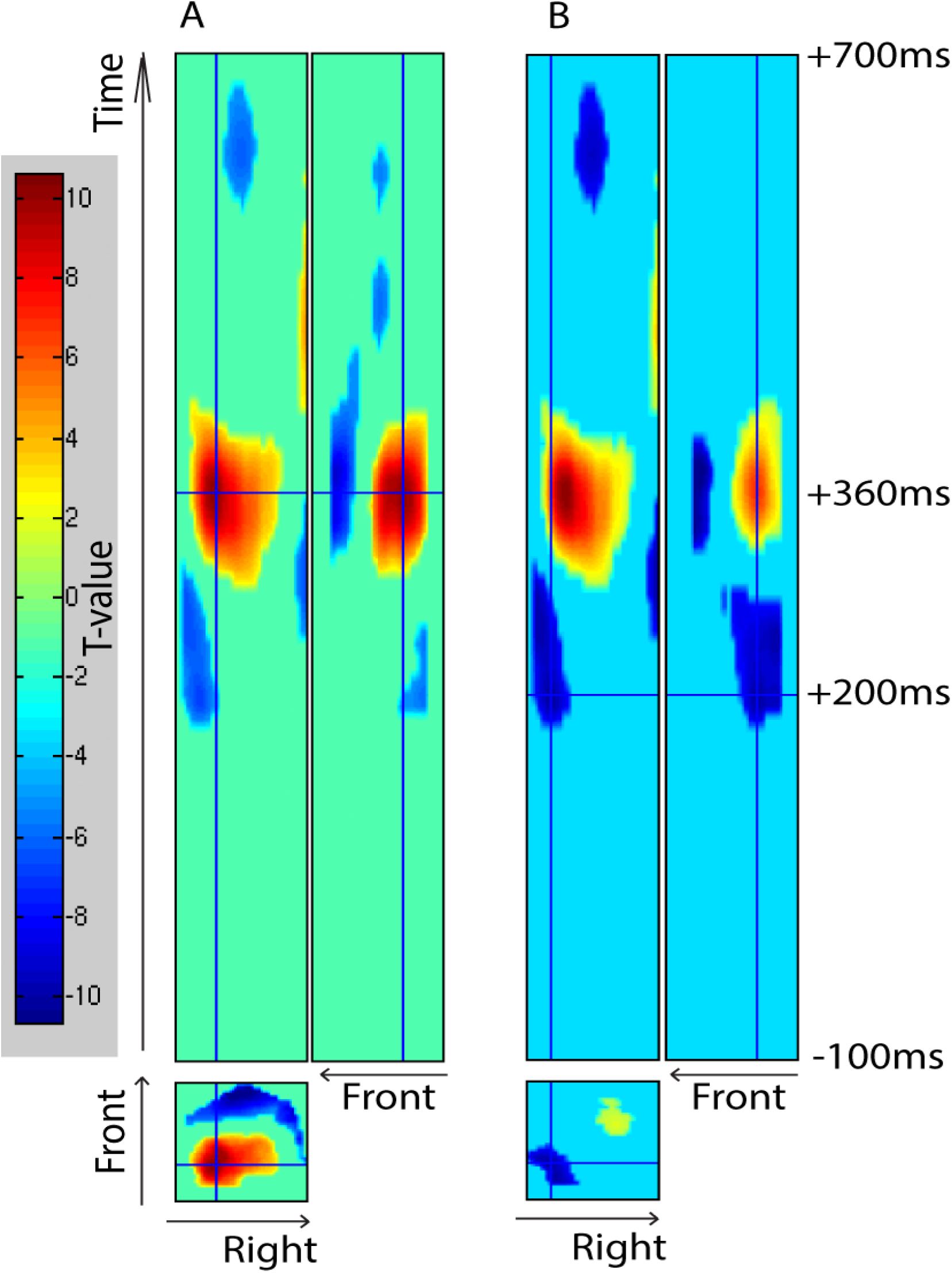
Sections through the 3D SPM for the ERP contrast of primed versus unprimed *same* trials in the same-different task. The sections in Panel A are through significant clusters that were positive over centroparietal sites (similar to Figure 3) for same judgments around 360ms, while the sections in Panel B are through clusters that were negative over left temporo-occipital sites around 200ms. See Figure 3 legend for more details.

**Figure 5:**
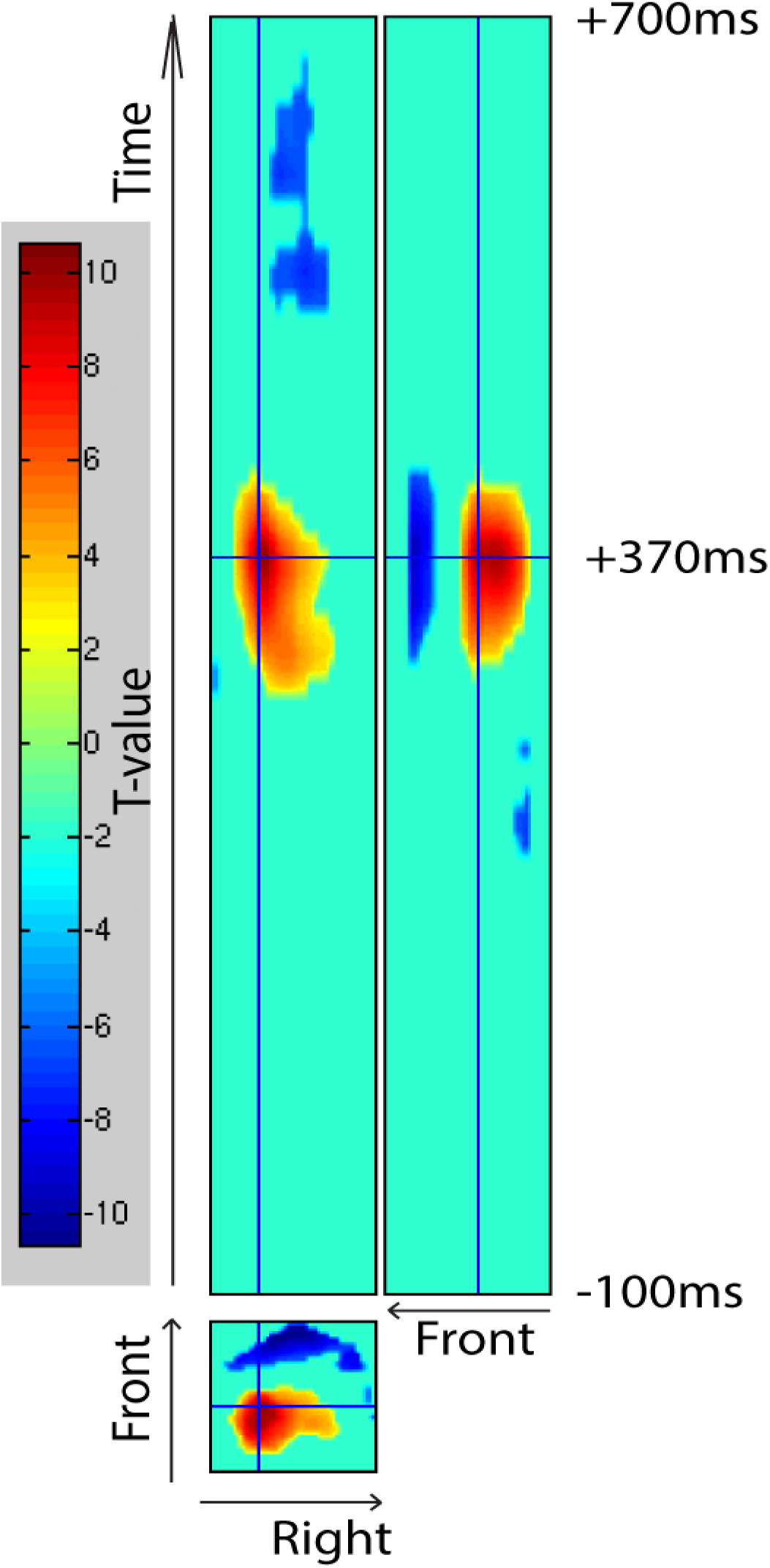
Sections through the 3D SPM for the ERP contrast of primed versus unprimed nonword *same* trials in the same-different task, through significant clusters that were positive over centroparietal sites (similar to Figure 4A) for same judgments around 370ms. See Figure 3 legend for more details.

#### Priming in the lexical decision task

For priming of words in lexical decision, three clusters survived correction (Figure 3). These clusters reflected a single topographic effect, comprising a central-parietal cluster that was more positive-going for primed relative to unprimed, accompanied by left and right fronto-temporal clusters of the opposite polarity, ranging between 360-540ms and peaking around 460ms. This most likely reflects a temporal shift of a central, positive-going component (as in Figure 2). No other word priming effects were found at other latencies. Nor did any clusters reach significance in the comparison of primed and unprimed nonwords. The interaction between priming and lexicality also peaked around 460ms.

#### Priming in the same-different task

As would be expected from the fact that the behavioural priming is greater in the same-different task than in lexical decision, the priming effects in the ERP data are also larger and onset earlier in time. For identity versus control *words* in *same* trials, a similar pattern of centroparietal positive-going differences accompanied by lateral negative-going differences was seen as for the lexical decision task (cf. Figure 4A and Figure 3), except that this priming effect became significant earlier, peaking around 360ms. Later, the polarity of this effect reversed, with a central negative-going difference from 450-620ms (peaking around 490ms) accompanied by fronto-lateral positive-going differences. This reversal most likely simply reflects a shift of a positive-going component earlier in time caused by priming (indeed, the same subsequent reversal of polarity was seen in the lexical decision task, but its extent did not reach significance owing to truncation by the end of the epoch).

The priming effects for nonwords in *same* trials were very similar to those for words, with a cluster from 280-420ms (peaking at 370ms) with more positive-going potentials over centroparietal sites for identity versus control nonwords, accompanied by fronto-lateral negative-going differences (Figure 5). The priming effects for words and nonwords are so similar that over the time range 280-420ms the waveforms for words and nonwords lie almost exactly on top of each other.

For *different* trials in the same-different task, no clusters reached significance for the comparison of same versus control trials, for either words or nonwords. Not surprisingly, the interaction between priming and same-different also peaked around 370ms.

For words in same trials there was an additional, earlier negative-going difference for identity versus control words over left temporo-occipital electrodes that survived correction from around 170-300ms (peaking around 200ms; Figure 4B). Interestingly, this effect also survived p<.001 in the SPMs for the primed vs unprimed comparison for words in the lexical decision task (visible in rightmost panel of Figure 3), though its extent did not survive correction for multiple comparisons. The same was also true for the primed vs unprimed comparison for different responses (visible in rightmost panel of Figure 5). Although not as robust as the later priming effects, this early ERP priming effect also patterns just like the behavioural priming.

Figure 6 shows the ERPs for each condition and task from electrode P9 that was closest to the statistical maximum of this early temporo-occipital priming effect. For lexical decision (Figure 6A), priming of words (but not nonwords) produces a more negative deflection of a N200 component, for both high and low frequency words (Figure 6B). For the same-different task, this early priming effect is even larger for both words and nonwords in same trials (Figure 6C) but was not significant for different trials (Figure 6D; see also Figure 7).

**Figure 6.**
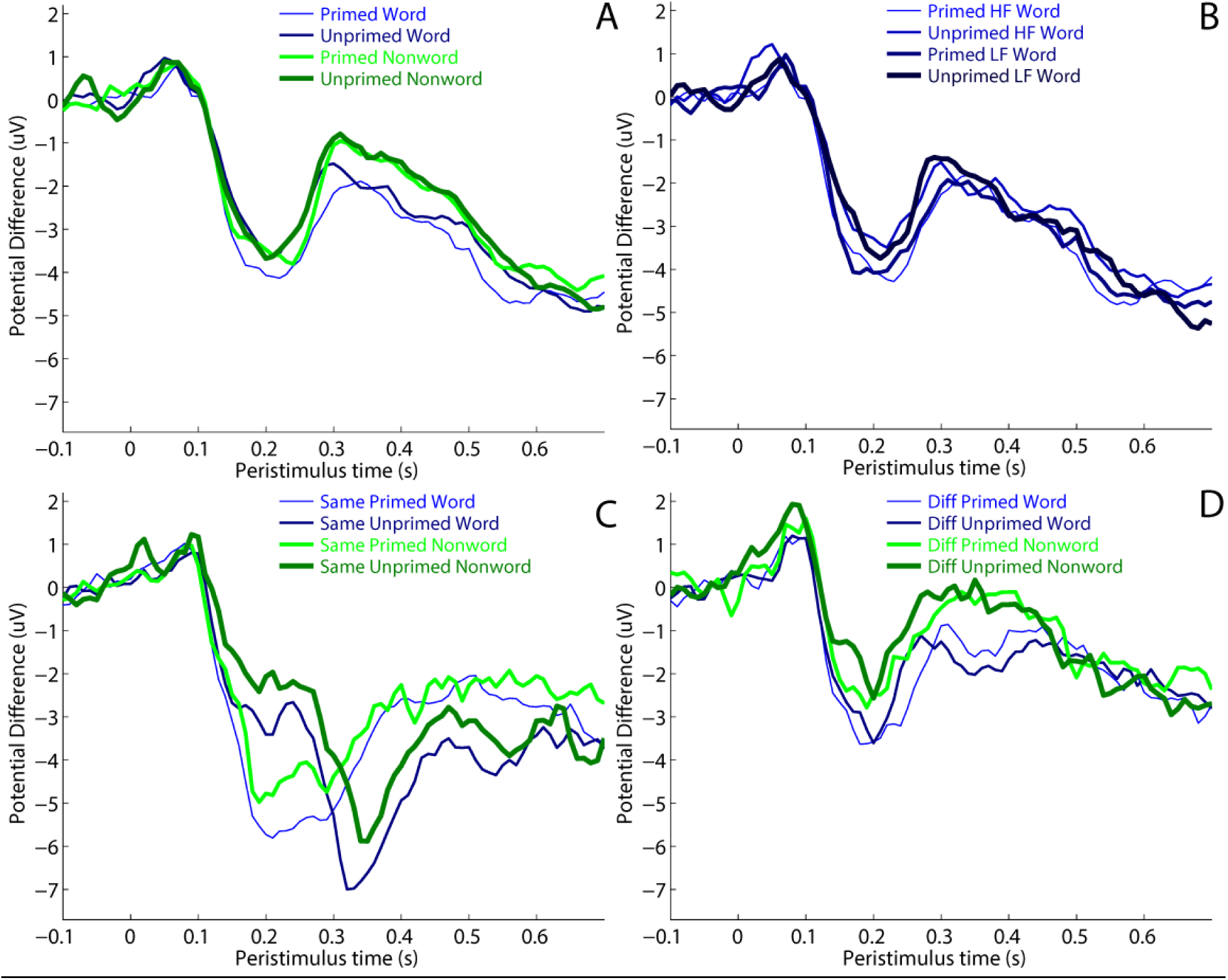
ERPs from temporo-occipital Channel P9 showing earlier priming effects for (A) primed and unprimed words and nonwords in lexical decision task, (B) primed and unprimed high frequency (HF) and low frequency (LF) words in lexical decision task, (C) primed and unprimed words and nonwords in same trials of same different task, and (D) primed and unprimed words and nonwords in different trials of same different task.

To summarise the effects of priming on the early, left temporo-occipital effect and later centro-parietal effect, we averaged the potential difference at these channels across 150-300ms and 300-500ms respectively. The results are shown in Figure 7, where the error bars represent 95% confidence intervals for the priming effect between consecutive conditions. Both effects showed similar patterns of priming across conditions, with the exception that priming for low-frequency words was significant in the early effect (Figure 7B) but did not reach significance in the later effect (Figure 7A). There was no evidence of significant differences (at this now uncorrected level of p<.05) for nonwords in lexical decision, or for words or nonwords in different trials of the same-different task.

**Figure 7.**
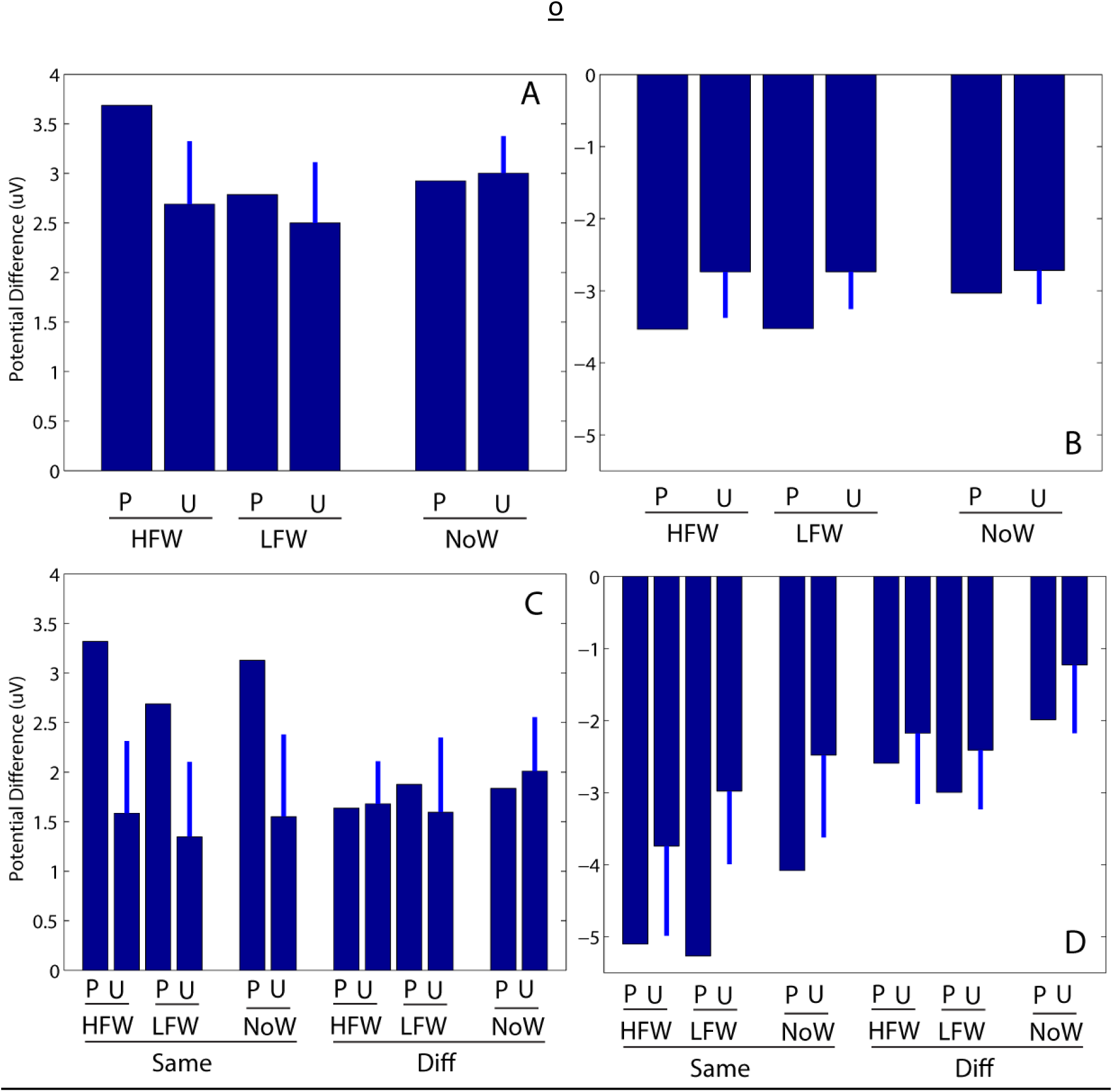
Average potential difference from 300-500ms for centro-parietal Channel P3 (Panels A+C) and from 150-300ms for temporo-occipital Channel P9 (Panels B+D) in lexical decision task (Panels A+B) and same-different task (Panels C+D). Error bars represent 95% confidence intervals for the priming effect between consecutive conditions on the x-axis. P = Primed; U = Unprimed; HFW = High Frequency Word; LFW = Low Frequency Word; NoW = Nonword.

### Statistical analysis of lexicality and frequency effects

In contrast to lexical decision, in the same-different task the effects of lexical status and word frequency on RTs are generally much weaker. In our same-different task there was a significant effect of lexical status which did not seem to be modulated by priming or response type (same vs. different), but no effect of word frequency. We find exactly the same pattern in the ERPs.

#### Lexical status in lexical decision

For primed words relative to primed nonwords in lexical decision, there was a central positive-going difference from 400-550ms (peaking at 480ms). This largely overlaps with the priming effect for words. It was accompanied by a smaller, later, and more posterior negative-going difference from 560-640ms (peaking at 610ms). There were no significant differences between words and nonwords for unprimed control trials.

#### Word frequency in lexical decision

In lexical decision, there was a central positive-going difference for high relative to low frequency words from 460-520ms (peaking at 490ms), which was significant for both primed and control conditions. It resembled the effect of lexicality in this task described above, even though the latter was only found for primed trials.

#### Lexical status in same-different

In the same-different task there was an overall effect of lexical status, averaged across *same* and *different* judgments, spread across three clusters: an early negative-going difference for words relative to non-words from 140-410ms over posterior left electrodes (peaking at 170ms;), followed by two positive-going clusters over central frontal electrodes from 240-340ms and 360-440ms (peaking at 280ms and 390ms respectively, but most likely part of the same effect). Note that the effect of lexical status was not significant when *same* and *different* responses were analysed separately.

#### Frequency in same-different

In the same-different task, there were no significant effects of word frequency for either same or different responses. Overall then, our two measures of lexical activation; word frequency and lexical status, appear different in the two tasks. Although there is an effect of lexical status in the same-different task, it is much weaker than in lexical decision and there is no sign of a frequency effect. This is the same pattern we see in the behavioural data.

#### Between task analysis

Any attempt to perform a combined analysis of the two task is complicated by the fact that, from the perspective of examining lexical effects, the structure of the two tasks is different. For example, words can only appear as words in lexical decision, but can appear in both the *same* and the *different* conditions in the same-different task. Also, given the complexity of the design, there are many different contrasts that could be examined. The most informative predictions that we can test are that word priming in lexical decision should be greater than the word priming in the *different* condition in the same-different task, and that the non-word priming in lexical decision should be smaller than the nonword priming in the same-different *same* condition. However, while the latter comparison revealed significant clusters around 340 and 370ms, no clusters were significant in the word comparison. This should not be too surprising. The nonword-same priming effect is very large relative to word priming in lexical decision. Indeed, word priming for that *same* condition is also bigger than word priming in lexical decision. However, given that the lexical effects in lexical decision are well established, the critical data for assessing the predictions of the Bayesian Reader come from the interactions between priming and *same* versus *different* responses, rather than from the comparison between experiments.

## Discussion

In both the behavioural literature and in electrophysiological studies, masked priming has frequently been promoted as providing insights into automatic processes involved in reading. In this context, ‘automatic’ has often been equated with ‘mandatory’, driven purely by the stimulus, irrespective of the task. In the electrophysiological literature, there has been the additional assumption that early EEG (and MEG) responses are a privileged index of automatic lexical processing, and therefore provide additional information beyond behavioural data. However, Norris and Kinoshita’s (2008) finding that the pattern of masked priming changes radically as a function of task calls these assumptions into question. According to the Bayesian Reader, priming is determined by the nature of the decision required by the task. In both lexical decision and same-different, the tasks require participants to judge whether the stimulus is a member of a particular set. Items in that set should show priming and items not in that set should not show priming. Here we have shown that not only are the behavioural results driven by the task, but the same is true of ERPs measured with EEG. The electrophysiological data mirror the behavioural data in that priming is obtained for *yes* responses in lexical decision and for *same* responses in the same-different task. Consequently in the same-different task it is possible to obtain priming for nonwords combined with an absence of priming for words. Reaction times in the same-different task are substantially faster than in the lexical decision task. This is also reflected in the ERP data, where waveforms in the same-different task are compressed and shifted forward in time. The strongest priming effects in the lexical decision and same-different tasks were found between 300-500ms, which would encompass the N400 component. Note that in the present context it is not important to attribute these effects to specific ERP components. However, masked priming has generally been assumed to modulate the N400 rather than, for example, the P3, and the N400 has been taken to index semantic or lexical processing (for reviews see Kutas et al. 2006; Lau et al. 2008; Grainger & Holcomb, 2009). Furthermore, the pattern of ERP effects we find in the lexical decision task is consistent with previous findings in the literature which have been assumed to reflect automatic lexical priming by masked stimuli. However, the fact that we see similar patterns in *same* responses to nonwords as in *yes* decisions to words in lexical decision shows that these ERP differences cannot be interpreted as being specifically lexical or semantic. We also found an early priming effect peaking at around 200 ms in both tasks. This effect almost certainly reflects early perceptual processes and is unlikely to be a consequence of any general speeding of motor responses in primed conditions.

More generally, the effect of priming seems better captured by an acceleration of the evoked response, rather than modulation of the amplitude of specific components (see also Henson, 2012). This general pattern is consistent with the Bayesian Reader account of masked priming. An identity prime acts to give a head start to target processing because the prime and target are effectively treated as the same perceptual event (see also Gomez, Perea & Ratcliff, 2013). An unrelated prime will cause interference, resulting in an overall slowing of target processing. It is this combination of facilitation and interference that makes it possible for the overall priming effect to sometimes be greater than the duration of the prime. The Bayesian Reader account also explains why this priming-related speeding of processing is greater in the same-different task than lexical decision task, as found here both in RTs and in the degree of acceleration of the ERPs (e.g, in terms of cross-over point).

In the introduction, we outlined three possible outcomes of this study: the electrophysiological data could track the behavioural data, they could remain the same despite the change in task, or the same-different task might simply overlay additional task-specific effects on top of automatic lexical effects. We can clearly rule out the second possibility; the task matters. The third possibility is harder to rule out definitively, as there may well be automatic lexical effects that we were unable to detect. Nevertheless, it does seem as though any residual automatic effects are very small, and certainly much smaller than the task-specific effects. This is hard to reconcile with Bowers’ (2010) suggestion that the pattern of priming in the same-different task is attributable to a lexical priming effect which is counteracted by a bias against responding the “Same” in the case of *different* words. (For behavioral evidence against the “bias” view, see Kinoshita & Norris, 2010, 2011). This view is also difficult to reconcile with the fact that the topography of the priming effect in lexical-decision and same-different is very similar. This suggests that they have a common mechanism and that priming in the same-different task is not the result of a combination of two opposing mechanisms.

Overall, the data are most consistent with the idea that the electrophysiological data exactly parallel the behavioural data. There is no indication that the same-different task elicits the same pattern of automatic ERP responses as in lexical decision, and that these are then overlaid with additional processes that operate only in the same-different task. At this point it is interesting to ask how our results might have been interpreted if we only had data for words in the same-different task. We think that the natural interpretation would have been that the same-different task magnifies the lexical priming effect. What implications do our results have for studies that have used masked priming to draw inferences about lexical processing? Although the pattern of results is task-specific, this does not necessarily imply that priming in lexical decision is in some sense ‘non-lexical’. As witnessed by the fact that priming is sensitive to lexical status and word frequency, priming clearly does reflect lexical knowledge. What these results do tell us is that *priming* is not an automatic consequence of reading a word – the nature of the decision process dynamically changes the relation between the prime and the target, and this is equally true of behavioural and electrophysiological responses. The change in the pattern of priming, in both behavioural and electrophysiological measures is a consequence of the change in the way the stimuli are processed in different tasks.

The current demonstration that electrophysiological measures can be dramatically modulated by changes in the nature of the task also has important implications for the design and interpretation of electrophysiological studies. First, we cannot assume that electrophysiological measures have a privileged status over behavioural measures in providing insights into early automatic processes. This might sometimes be the case, but we cannot make such an assumption without first demonstrating task independence. However, the most important point is a very simple one; the task matters. The nature of the task determines the configuration of both the cognitive and the neural processes that are engaged.

## Appendix

### High-frequency word targets

club, home, well, team, army, walk, room, size, park, love, hall, felt, sign, case, part, fire, time, hand, late, news, side, cold, trip, data, body, wife, farm, step, work, full, name, over, rose, hair, free, town, book, good, ball, thin, child, scene, young, sense, brown, table, south, class, press, blood, house, month, think, black, broad, least, clear, share, green, plant, leave, women, water, group, major, death, white, early, heavy, drive, hotel, earth, story, heart, after, money, peace, voice, trade, large, field, trial, river, price, level, might, round, north, night, could, shall, stock, small, while, wrong, party, along, stood, basic, years, plane, daily, court, today, still, woman, below, reach, stage, value, order, every, great, human, issue, first, among, those, ready, under, bottom, simple, office, island, twenty, course, public, strong, degree, health, mother, toward, really, number, spring, church, reason, answer, couple, window, before, inside, social, enough, letter, person, nature, theory, battle, income, friend, school, market, centre, figure, united, pretty, common, family, street

### Low-frequency word targets

ache, jeep, mint, doll, limb, zone, exit, quiz, fowl, sofa, bake, scar, toil, hawk, pony, lung, cage, wink, knit, flex, dumb, rope, arid, gash, oven, hunt, arch, kiss, lamp, vase, flip, rail, numb, gown, tank, bald, peak, idle, mist, sail, thigh, plank, brick, blast, squat, hound, towel, greed, blush, bough, sonar, blink, adopt, crest, bland, creep, stare, steer, creed, chore, wheat, lemon, scout, flour, grill, glove, petty, boast, arrow, agony, super, knock, bleak, brisk, frail, pearl, spade, atlas, wreck, demon, trunk, toast, lyric, comic, canoe, filly, sleet, greed, minus, roast, baton, sewer, timer, slack, repel, crepe, scare, cheek, slats, steak, niece, essay, attic, thief, barge, lunar, purse, sport, sweep, blaze, feast, ankle, globe, gland, virus, candy, cargo, mouse, booth, weird, barrow, quarry, uproar, grocer, peanut, litter, vendor, hazard, drawer, poison, opaque, hammer, rescue, nugget, willow, enamel, toffee, import, copper, hollow, sailor, guitar, needle, canyon, launch, ransom, jockey, bronze, marina, coyote, occupy, invade, jungle, shrine, enigma, fabric, wallet, morale, kidney, tennis

### Nonword targets

dern, dirp, blit, yebb, stib, yalt, bymn, nurf, mawk, yarm, sarp, yurk, jark, gube, flis, zere, jipe, yeap, cilm, bleg, glon, ferg, snav, jush, filk, flun, zean, voun, nurl, flub, yiek, zodd, vaif, tirp, hulf, yirt, jisp, nuip, rurp, norb, cloor, shend, shice, breet, thare, frone, beark, truld, prout, gound, wouch, trand, cherd, sterm, thace, theak, whert, thirp, dreat, breat, blout, sonth, graim, pight, selch, minch, fleep, starp, thest, rilse, chesk, hilch, thove, drash, slart, frind, smace, blace, bline, theen, trong, theng, nould, chout, grald, coulk, sould, grich, friss, roult, flost, wonth, shong, whosk, clest, courm, thean, shair, smair, plart, shisk, chand, blong, chall, stire, banch, whoce, brult, chist, litch, pring, dring, therp, sterk, sping, shint, woult, glast, preat, broul, thrump, phloof, cherse, stonte, frutch, cheash, thouth, slorse, slitch, phrull, snorge, knooch, spreep, flenge, posque, blourn, thulge, stralt, thwirk, thrurp, sploan, thwoop, gwodge, thorth, splame, skoosh, proque, dweeth, plynch, sprobe, glouge, scidge, cleech, florge, shramp, vauche, cheath, chench, brange, sploap

### High-frequency word control primes

heat, last, back, long, once, hour, bank, land, feet, dark, neck, wide, real, unit, mind, both, word, rest, form, high, turn, life, show, live, test, took, blue, clay, deep, stop, lost, wait, fact, open, pain, area, idea, year, very, lord, range, total, where, local, state, force, piece, teeth, final, image, again, serve, power, front, seven, above, often, began, start, write, third, until, union, chief, spent, floor, march, thing, point, about, bring, sound, given, music, which, built, truth, staff, eight, visit, start, seven, about, staff, point, speak, visit, image, place, serve, write, began, eight, truth, cause, sound, music, piece, teeth, built, chief, spent, given, where, range, third, march, until, floor, bring, again, total, union, power, front, above, state, final, thing, local, energy, county, actual, though, should, taking, method, always, action, bridge, annual, single, moment, attack, cattle, system, active, second, bright, caught, within, doctor, extent, status, famous, middle, simply, making, crisis, hardly, almost, indeed, supply, follow, cannot, across, fiscal, weight, ground, policy

### Low-frequency word control primes

stud, omit, bell, grip, tune, acid, wool, wand, bunk, hump, wolf, monk, rump, fuel, zest, heir, lush, toss, grab, oath, weep, chum, pulp, mock, suck, boil, link, germ, coin, plum, vest, bush, dish, bark, weed, prey, cult, tart, glow, nude, abbey, sober, fever, swell, flock, stack, scrap, arena, stamp, lever, juice, torso, nurse, gloom, choke, float, chill, cliff, blunt, bland, slump, sword, alarm, width, couch, bunch, choir, ivory, shelf, exert, alibi, medal, ghost, patch, dough, onion, thumb, merry, salad, quack, salad, merry, quack, fever, drift, torso, onion, stamp, dwell, juice, nurse, gloom, flock, dough, stack, blunt, thumb, float, choke, brood, sword, print, shelf, alarm, slump, exert, width, medal, choir, ghost, ivory, couch, patch, sober, abbey, lever, swell, alibi, arena, bunch, giggle, knight, helium, plasma, sorrow, banana, alight, nibble, chubby, breeze, clinic, clutch, lagoon, raisin, parent, bishop, quaint, bauble, alumni, scream, dented, wobble, tomato, esteem, exempt, velvet, wintry, addict, locker, salami, senile, gloomy, potato, mammal, scotch, solemn, circus, filthy, saloon, orphan

### Nonword control primes

phim, noak, nawk, vauk, glof, curn, zice, tisp, frut, cheg, cluy, bobe, yome, vant, bebb, droy, nazz, swoz, goaf, phav, cemp, julk, kerp, cimb, gwob, spro, smor, drek, yemp, nawn, nalf, frap^*^, pham, gowl, jese, mune, darb, coth, jisk, suzz, deash, bruck, broon, shool, clond, chilk, thind, sheck, sheel, chirk, trape, fluge, slass, chond, wrunk, lorch, sland, lough^*^, floop, shund, chade, quile, clost, warld, frint, wreat, torth, glome, pronk, fooph, glain, bedge, crawn, flide, dench, hathe, brith, shird, crame, crulb, sheel, fooph, trape, fluge, theck, brate, brith, sland, clond, deash, bruck, shird, prike, thand, broon, deast, lorch, wrunt, lench, shend, wreat, quile, chade, torth, warld, glome, frint, flide, pronk, crawn, clost, hathe, shool, glain, crulb, crame, bedge, chirk, chilk, steck, splegg, swudge, ploint, chanch, splasp, fluint, frenge, dwunch, grodge, gwadge, plawre, squalm, chouch, dratch, wrilch, swetch, sproon, gnunge, clunge, thrimp, guiche, scrull, plutch, glauve, floach, draunt, thwail, smuick, skidge, dwaunt, straif, throll, skount, thwick, dwulge, sploob, splink, spladd, thoosh, thuiff

### High-frequency words; different referents

door, born, face, girl, hold, look, road, city, wish, bill, food, paid, king, down, mass, feed, near, nine, post, fell, block, bread, cable, clean, dress, moral, glass, greet, flood, lease, space, radio, paper, plans, horse, sharp, mouth, thank, youth, three, doubt, basis, shell, smell, faith, parts, alone, light, right, bound, yearn, forth, found, stick, stool, study, whole, would, wring, short, growth, amount, little, result, modern, volume, summer, police, dinner, normal, winter, corner, effort, spirit, nation, future, father, square, report, design

### Low-frequency words; different referents

curd, flee, gaze, moss, bail, comb, knob, heap, pole, ruin, bang, knot, rake, itch, bump, quit, swim, hook, pear, tomb, chord, algae, breed, creek, adapt, stark, cough, steep, blank, crust, dodge, flush, logic, mound, polar, scrub, creep, blond, solar, vowel, alley, bacon, boast, slick, sever, slate, crept, rebel, elbow, silly, scars, fleet, shaft, cheer, quill, sinus, freed, steam, tenth, tiger, salmon, shrink, ambush, blouse, gender, cheese, dental, legacy, mellow, cavity, facial, escort, clergy, dinghy, mildew, blonde, humane, nutmeg, borrow, superb

### Nonwords different referents

thim, sich, tuip, koof, twag, ghez, dwom, wrid, spee, kont, horb, sneb, wudd, hube, mimb, whad, noof, relk, blod, faph, trind, bruld, cousk, thear, brean, geark, moung, dreak, starm, chird, thice, brone, shent, cound, thire, carth, drout, mousk, wherp, worch, bleam, boult, shain, briss, moulk, shing, thead, couse, jance, flast, skirm, shace, coure, brich, yould, plard, smain, whesk, wouth, clast, bleant, thruke, shrobe, phrass, thrunt, splawn, phanch, platch, speeth, geethe, pheash, brulch, phross, phramp, broosh, thwegg, plarce, dwogue, strisk, crowth

## Footnotes

1 It is important to emphasize that what is integrated is the evidence used to make the decision, not the low-level perceptual data (e.g., visual features in the prime and the target), as attested by the fact that masked priming effects in visual word recognition tasks are independent of the visual similarity of the lowercase prime and uppercase target (Bowers, Vigliocco, & Haan, 1998; Kinoshita & Norris, 2008).

2 This is apparent in Figure 2C where priming leads to an increase in the amplitude between about 250-400ms, but a decrease thereafter. A window centred around 400ms might show little effect of priming. See Kilner (2013) for further problems with choice of time windows.

* Although rare, ‘frap’ and ‘lough’ are both words. However, they were not part of the analysis as they were control primes only.

